# SpliceRead: Improving Canonical and Non-Canonical Splice Site Prediction with Residual Blocks and Synthetic Data Augmentation

**DOI:** 10.64898/2026.02.05.703825

**Authors:** Sahil Thapa, Khushali Samderiya, Rohit Menon, Oluwatosin Oluwadare

**Affiliations:** Department of Computer Science and Engineering, University of North Texas, Denton, 76203, Texas, USA; Center for Computational Life Sciences, University of North Texas, Denton, 76207, Texas, USA; Department of Computer Science, University of Colorado at Colorado Springs, Colorado Springs, 80918, Colorado, USA

**Keywords:** Splice site prediction, Non-canonical sequences, Transcriptomic, Deep learning, Residual CNN, Data augmentation, ADASYN, Genomic classification, Cross-species generalization

## Abstract

Accurate splice site prediction is fundamental to understanding gene expression and its associated disorders. However, most existing models are biased toward frequent canonical sites, limiting their ability to detect rare but biologically important non-canonical variants. These models often rely heavily on large, imbalanced datasets that fail to capture the sequence diversity of non-canonical sites, leading to high false-negative rates. Here, we present SpliceRead, a novel deep learning model designed to improve the classification of both canonical and non-canonical splice sites using a combination of residual convolutional blocks and synthetic data augmentation. SpliceRead employs a data augmentation method to generate diverse non-canonical sequences and uses residual connections to enhance gradient flow and capture subtle genomic features. Trained and tested on a multi-species dataset of 400- and 600-nucleotide sequences, SpliceRead consistently outperforms state-of-the-art models across all key metrics, including F1-score, accuracy, precision, and recall. Notably, it achieves a substantially lower non-canonical misclassification rate than baseline methods. Extensive evaluations, including cross-validation, cross-species testing, and input-length generalization, confirm its robustness and adaptability. SpliceRead offers a powerful, generalizable framework for splice site prediction, particularly in challenging, low-frequency sequence scenarios, and paves the way for more accurate gene annotation in both model and non-model organisms.The open sourced code of SpliceRead and a detailed documentation is available at The open-sourced code of SpliceRead and detailed documentation are available at https://github.com/OluwadareLab/SpliceRead.

## 1 Background

Splice sites are crucial regions within pre-mRNA that define the boundaries of exons and introns, enabling accurate splicing during the gene expression process [1, 2]. These sites are broadly categorized into *acceptor* and *donor* types, representing the 3’ and 5’ ends of introns, respectively [3, 4]. Acceptor sites typically mark the beginning of an exon and are denoted by the AG sequence, while donor sites denote the end f an exon and are denoted by the GT sequence [1, 2, 5]. Accurate identification of both types is critical for understanding the complete splicing mechanism, as errors in splicing can lead to severe genetic disorders and diseases [3, 6, 7].

Splice sites are further divided into *canonical*, which conform to the consensus sequences (e.g., AG and GT), and *non-canonical*, which deviate from these sequences [5, 8]. Non-canonical acceptor and donor sites, in particular, exhibit diverse sequence patterns, making them harder to classify and demanding innovative computational approaches [3, 4, 9]. Recent advancements in machine learning and deep learning have revolutionized splice site prediction. Convolutional Neural Networks (CNNs), in particular, have shown great promise in identifying splice sites with high accuracy [1– 4, 6, 10–12]. Transformers and large-scale genomic models have also entered the field, with tools such as DNABERT [13] and the Nucleotide Transformer [14] showcasing cross-domain sequence learning capabilities.

Building on these advancements, researchers have proposed various deep learning models to address key challenges. Wang et al. [1] introduced *SpliceFinder*, an ab initio CNN-based model for splice site detection, showing improved accuracy over traditional methods like MaxEntScan [8] and NNSplice [15]. Jaganathan et al. [9] proposed *SpliceAI*, a deep residual neural network that models long-range sequence context to predict splice donor and acceptor sites directly from primary genomic sequence. Scalzitti et al. [2] developed *Spliceator*, a multi-species predictor leveraging CNNs for generalization across organisms. Despite their success, these models often struggled with imbalanced datasets and limited non-canonical performance. To improve robustness, Akpokiro et al. [3] proposed *CNNSplice* with advanced preprocessing techniques, while Akpokiro et al. [6] introduced *EnsembleSplice*, combining multiple architectures for better accuracy. More recently, Chao et al. [4] presented *Splam*, a deep-learning-based predictor designed to improve spliced alignments. However, its alignment-focused training limits its utility in non-canonical splice site prediction. Newer genome-wide tools like MapSplice [16], TopHat [17], and StringTie [18] further emphasize alignment accuracy but do not explicitly target non-canonical classification.

Hence, in this study, we build on prior work by addressing these limitations through a novel deep learning model—SpliceRead—that integrates residual blocks [19] and synthetic data augmentation to improve performance, particularly for non-canonical splice sites. We incorporate synthetic oversampling via ADASYN [20] to correct class imbalance, and use batch normalization [21] and dropout for training stability. Our evaluation strategy relies on comprehensive metrics as recommended by Sokolova and Lapalme [22], Powers [23], and we apply SHAP for interpretability [24].

This paper aims to:

- Improve multiclass classification of both canonical and non-canonical splice sites via a novel CNN architecture with residual blocks.
- Address class imbalance through synthetic data augmentation [20].
- Evaluate performance across splice site subtypes using robust evaluation metrics [22, 23].
- Demonstrate the benefits of synthetic augmentation in biological sequence learning.
- Validate cross-species performance to assess generalizability and adaptability across diverse genomes.

## 2 Materials and Methods

### 2.1 Dataset

The datasets used in this study were sourced from Spliceator, a comprehensive resource for splice site prediction [2]. These datasets were constructed using sequences rom the G3PO+ dataset, which includes 2,741 sequences from 147 phylogenetically diverse organisms [2]. Each sequence is classified as either ‘Confirmed’ (error-free, 1,361 sequences) or ‘Unconfirmed’ (containing gene prediction errors, 1,380 sequences). To ensure high-quality data, only ‘Confirmed’ sequences were used to construct the Gold Standard (GS) dataset [2].

For this study, GS was selected as the positive subset, containing 10,973 donor and 11,179 acceptor splice site (SS) sequences. Sequences were standardized to two lengths: 400 and 600 nucleotides, ensuring that splice site classification was robust across different input sizes [2]. The splice site is centrally positioned at bases 201 and 202 for 400-length sequences and 301 and 302 for 600-length sequences, marked by the canonical “GT” (donor) or “AG” (acceptor) dinucleotide. Any sequences containing undetermined nucleotides (“N”) or duplicates were removed to maintain data integrity.

The negative subset was selected from GS 1, which comprises sequences that lack splice sites, ensuring a controlled negative dataset for training and evaluation. Table 1 provides a summary of the dataset composition, including the number of canonical and non-canonical sequences for positive subsets and the total number of negative sequences used in this study.

**Table 1:** Overview of dataset composition used in this study. The positive subset includes confirmed canonical and non-canonical splice sites from the GS1 dataset. The negative subset comprises sequences without splice sites, used as controls. Sequences cover both acceptor and donor sites, with lengths of 400 and 600 nucleotides to support robust prediction.

| Dataset | Canonical | Non-Canonical | Negative |
| --- | --- | --- | --- |
| Acceptor | 11,036 | 143 | 11,000 |
| Donor | 10,736 | 237 | 11,000 |
| **Total** | **21,772** | **380** | **22,000** |

All sequences were organized within a structured directory format, ensuring systematic storage of positive and negative subsets while facilitating efficient data processing for training and evaluation. The inclusion of both 400 and 600 nucleotide sequence lengths ensures that the model can generalize across different input sizes, enhancing splice site prediction robustness. These datasets form the foundation for assessing splice site classification performance, particularly in distinguishing canonical and non-canonical splice sites [2].

### 2.2 Data Preprocessing

For efficient processing of genomic sequences, we applied one-hot encoding, a standard technique in deep learning for DNA sequence analysis, converting each nucleotide into a binary vector of size four: adenine (A) as (1, 0, 0, 0), cytosine (C) as (0, 1, 0, 0), guanine (G) as (0, 0, 1, 0), and thymine (T) as (0, 0, 0, 1). Indeterminate nucleotides (‘N’) were encoded as (0, 0, 0, 0) to minimize ambiguity [2]. Each encoded sequence was represented as a first-order tensor of length *W* (*W* = 600 or 400), and the complete dataset formed a four-dimensional tensor of shape *S* × *H* × *W* × *C*, where *S* is the number of sequences, *H* = 1 the height of the 1D vector, *W* the sequence length, nd *C* = 4 the nucleotide channels [2]. Sequences containing excessive ‘N’ bases were excluded to ensure data integrity, standardizing inputs and enabling convolutional ayers to effectively extract spatial dependencies relevant to splice site prediction.

### 2.3 SpliceRead Model Architecture

SpliceRead is a deep convolutional neural network (CNN) designed for accurate splice site classification. The model operates on one-hot encoded DNA sequences and employs residual connections to enhance feature extraction while mitigating vanishing gradient issues. The complete architecture consists of four main stages: input representation, data augmentation, deep learning block, and classification (see Figure 1).

**Fig. 1:**
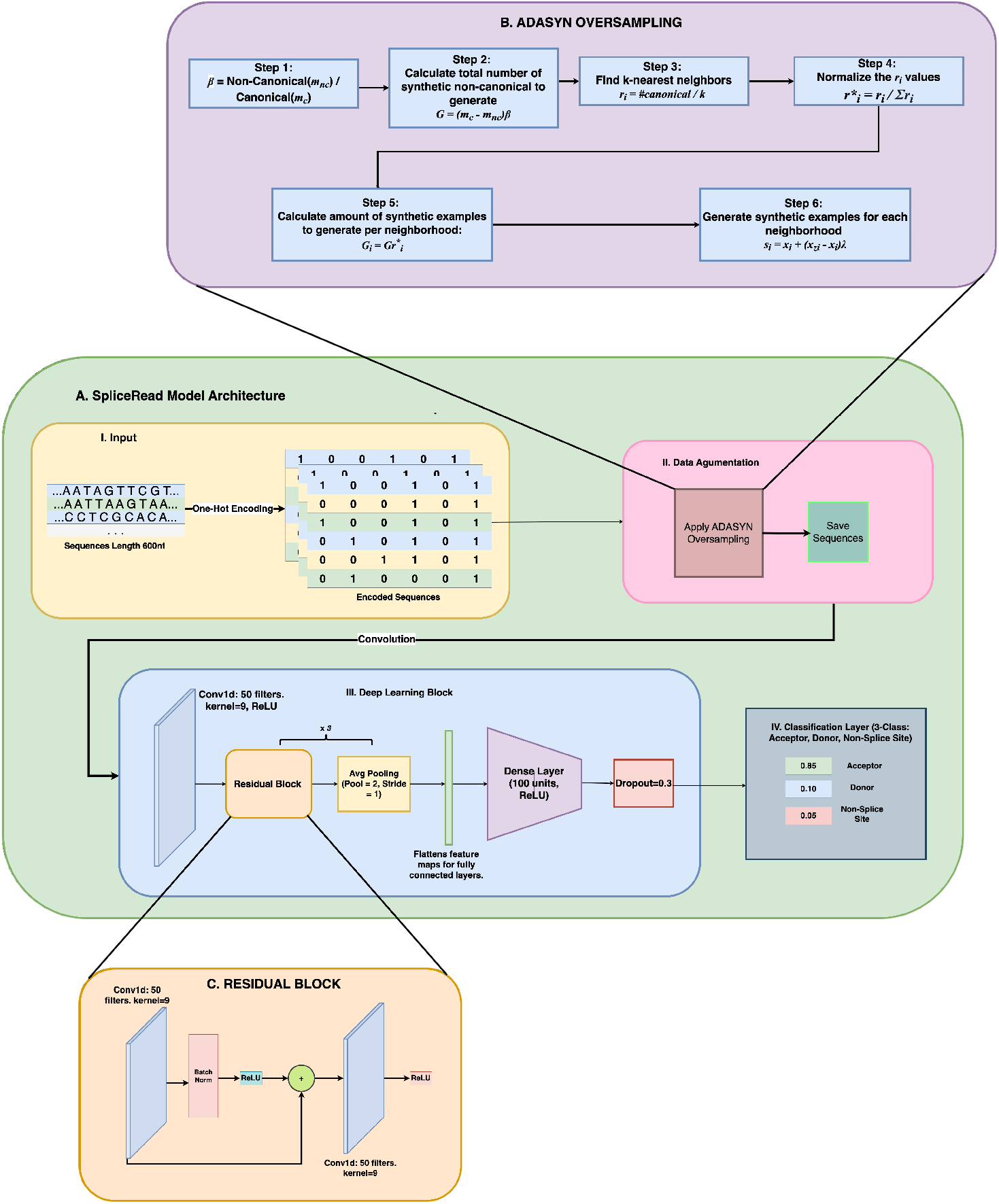
SpliceRead Model Architecture. The model consists of four major blocks: (**A**) *Input Block* (**B**) *Data Augmentation Block*, (**C**) *Deep Learning Block* (**D**) *Classification Block* This architecture helps capture local dependencies, balance class distribution, and improve performance on rare non-canonical splice sites.

#### 2.3.1 Input Representation

The model takes genomic sequences of fixed lengths (400 or 600 nucleotides) as input. Each nucleotide in the sequence is converted into a one-hot encoded representation, resulting in matrices of shape 400×4 or 600×4, where each column corresponds to one of the four nucleotides: adenine (A), cytosine (C), guanine (G), and thymine (T) [1]. his encoding ensures that spatial dependencies between nucleotides are preserved, allowing the convolutional layers to learn meaningful sequence features (see Figure 1-A. I. Input).

#### 2.3.2 Data Augmentation

One of the major challenges in splice site prediction is the severe imbalance between canonical and non-canonical sequences, where canonical sites vastly outnumber non-canonical ones, often leading to biased models that overfit to the dominant class. This imbalance affects the accurate prediction of non-canonical sites. To mitigate his issue, we explored two augmentation methods, including Borderline Synthetic Minority Over-sampling Technique (Borderline-SMOTE) [25] and Adaptive Synthetic Sampling (ADASYN) [26]. Among these, ADASYN [26] produced the best results and was adopted as our augmentation strategy. ADASYN generates synthetic samples by interpolating between minority-class instances—in this case, non-canonical splice sites—creating augmented datasets with varying synthetic-to-real data ratios (100%, 80%, 60%, 40%, 20%, 15%, 10%, and 5%). The approach reduces class imbalance and improves generalization by expanding the feature space of rare non-canonical patterns, exposing the model to a broader range of biologically relevant variations.

To ensure effective learning, the data were first divided into canonical and non-canonical subsets, and ADASYN interpolated new samples within the minority-class feature space until the desired target count was achieved. The generated one-hot vectors were decoded back into nucleotide sequences using inverse transformation to form multiple augmented datasets. For each augmentation ratio, the number of synthetic non-canonical sequences was computed as: *Synthetic Count* = [Ratio (%) × *Number of Canonical Sequences*] − *Number of Real Non-Canonical Sequences*. For example, in the 100% setting, synthetic sequences matched the canonical count, whereas in the 80% setting, 80% of the canonical count defined the target, and the remainder deter-mined the synthetic count. This procedure was repeated for all ratios down to 5%. Based on the results presented in the *Results* section, a 5% synthetic-to-real ratio was selected as the default configuration, with the augmented sequences integrated into the training pipeline alongside real data (see Figure 1A–II).

#### 2.3.3 Deep learning Block

To enhance feature extraction and gradient flow, *SpliceRead* employs three stacked residual blocks, each containing two 1D convolutional layers with 50 filters and a kernel size of 9, followed by batch normalization [21] and ReLU activation. Skip connections link each block’s input to its output, mitigating vanishing gradients and enabling identity mapping [19]. Average pooling (pool size = 2, stride = 1) between blocks downsamples feature maps while retaining biologically relevant patterns, allowing capture of both short- and long-range dependencies. The final block output is flattened and passed through a dense layer with 100 ReLU units, followed by dropout (rate = 0.3) and a softmax layer classifying acceptor, donor, and non-splice sites. The model is trained with Adam (learning rate 1 × 10^−4^) using categorical cross-entropy loss and evaluated via five-fold cross-validation, achieving robust classification performance, particularly for rare non-canonical splice sites (Figure 1A–III).

#### 2.3.4 Classification Block

The output features from the final residual block are flattened and passed through a dense layer with 100 ReLU-activated neurons. A dropout layer (rate = 0.3) is applied to reduce overfitting. Finally, a softmax layer outputs class probabilities for three categories: acceptor site, donor site, and non-splice site. This enables the model to perform multi-class classification of genomic sequences based on splice site type. (Figure 1-A. IV. Classification Block).

### 2.4 Training and Testing Setup

The model was trained and evaluated using a deep CNN with residual blocks to classify splice sites into four categories: canonical and non-canonical acceptors and donors. The dataset was split into 80% for training/validation and 20% for testing, with the test set kept completely unseen during development to prevent overfitting. Five-fold cross-validation was employed, where each fold trained on four subsets and validated on the fifth, ensuring robust generalization. The model was optimized using Adam (learning rate 10^−4^) with categorical cross-entropy loss and trained using mixed-precision under CUDA 12.0 for computational efficiency.Checkpointing preserved the best-performing model based on validation accuracy. Training and testing were performed on an Ubuntu 24.03.3 LTS workstation with an NVIDIA GeForce RTX 4090 GPU (24 GB VRAM), Intel(R) Xeon(R) w7-3445 CPU, and 125 GB RAM. Implemented in TensorFlow with GPU acceleration, each trained model was saved for structured comparisons, and the best model from cross-validation was finally evaluated on the isolated test set to provide an unbiased estimate of real-world splice site classification performance.

### 2.5 Evaluation Metrics

To evaluate model performance, we used standard classification metrics—accuracy, recision, recall, and F1-score [22, 23]—which collectively assess prediction quality, especially in imbalanced datasets where minimizing both false positives (FP) and false negatives(FN) is crucial. In addition, we introduced two misclassification metrics to assess how effectively the model distinguishes canonical and non-canonical splice sites. The Non-Canonical Misclassification Rate (NCMR; Equation 1) measures errors pecific to non-canonical sites:

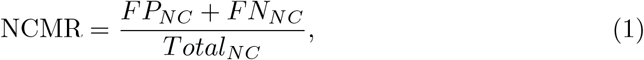

where *FP*_*NC*_ and *FN*_*NC*_ are the counts of non-canonical sites misclassified as canonical or non-splice sites, and *Total*_*NC*_ is the total number of non-canonical sites. Similarly, the Canonical Misclassification Rate (CMR; Equation 2) quantifies misclassification of canonical sites:

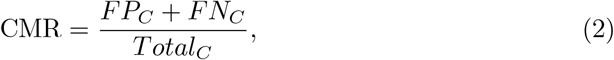

where *FP*_*C*_ and *FN*_*C*_ denote canonical sites misclassified as non-canonical or non-splice sites, and *Total*_*C*_ is the total canonical count. Lower NCMR and CMR values indicate improved discrimination between splice site types. Together, these metrics complement standard evaluations, offering a more nuanced and reliable assessment of the model’s generalization and accuracy in identifying both canonical and non-canonical splice sites.

## 3 Results

### 3.1 Hyperparameter Optimization

To achieve optimal performance in classifying canonical and non-canonical splice sites, careful tuning of model hyperparameters was necessary. The optimization process involved extensive experimentation with various architectural components, including the configuration of residual blocks and the parameters for data augmentation. These decisions were driven by the need to balance accurate feature extraction with generalizability across imbalanced genomic datasets.

#### Deep Learning Block Design and Hyperparameter Selection

The core architecture of *SpliceRead* employs residual connections to enhance deep feature learning and prevent vanishing gradients. Each residual block contains two 1D convolutional layers with batch normalization and ReLU activation. Through extensive experimentation, we selected three residual blocks, which balanced model depth and overfitting; 50 filters per convolutional layer, capturing diverse local sequence patterns without redundancy; and a kernel size of 9, sufficient for short-to mid-range nucleotide dependencies. ReLU served as the activation function for its efficiency and stability, and average pooling (pool size = 2, stride = 1) preserved subtle spatial information essential for distinguishing non-canonical splice sites. These hyperparameters were optimized via five-fold cross-validation and chosen based on the highest F1-score across canonical and non-canonical splice site predictions, with the final configuration summarized in Supplementary Table S1.

#### Data Augmentation: Hyperparameter search

Augmented datasets were evaluated independently to determine the optimal level of augmentation, with final model selection based on the configuration that yielded the best overall classification performance across splice site subtypes. It is important to note that the total number of training samples varied across augmentation levels, as each setting introduced a different number of synthetic non-canonical sequences based on the target ratio. Table 2 provides an overview of the number of synthetic sequences generated across different augmentation levels. The datasets were systematically structured to facilitate effective model training and evaluation.

**Table 2:**
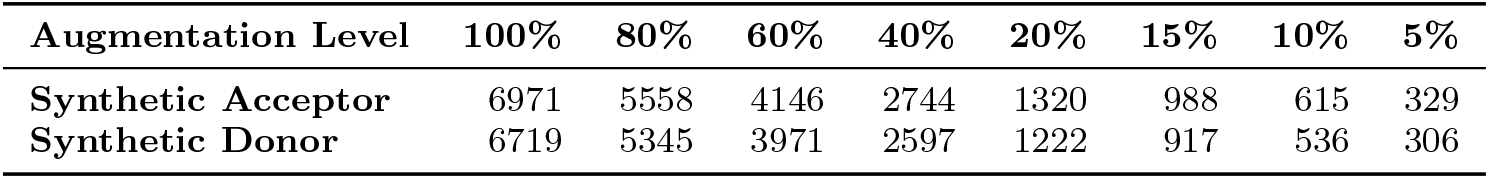
Synthetic non-canonical splice site generation using ADASYN at varying augmentation levels. This table shows the number of synthetic acceptor and donor sequences created for each augmentation level (100% to 5%).

| Augmentation Level | 100% | 80% | 60% | 40% | 20% | 15% | 10% | 5% |
| --- | --- | --- | --- | --- | --- | --- | --- | --- |
| **Synthetic Acceptor** | 6971 | 5558 | 4146 | 2744 | 1320 | 988 | 615 | 329 |
| **Synthetic Donor** | 6719 | 5345 | 3971 | 2597 | 1222 | 917 | 536 | 306 |

To visualize the distributional differences between canonical, non-canonical, and synthetic sequences, we plotted their GC and AT content distributions from (10 to 100% synthetic ratio). Supplementary Figures S1 and S2 illustrate these distributions. To evaluate the impact of synthetic data augmentation on splice site prediction, we trained the SpliceRead model using multiple datasets augmented with ADASYN at varying levels: 100%, 80%, 60%, 40%, 20%, 15%, 10%, 5%, and 0% synthetic-to-real data ratios. Each dataset was independently used to train the model via five-fold cross-validation to ensure statistical robustness.

Table 3 and Table 4 summarize the classification performance—measured by F1-score, CMR Score, and NCMR Score across each augmented dataset for both Donor and Acceptor sequences. The model trained without any synthetic data serves as a baseline for comparison.

**Table 3:** Evaluation of SpliceRead across ADASYN augmentation levels for Acceptor. This table reports F1 Score *±* Standard Deviation (SD), Acceptor CMR *±* Standard Deviation (SD), and Acceptor NCMR *±* Standard Deviation (SD) across folds for models trained with increasing levels of synthetic non-canonical data.

| Augmentation Level | F1 Score | Acceptor CMR | Acceptor NCMR |
| --- | --- | --- | --- |
| 0 | 0.9503 ± 0.0012 | 0.0528 ± 0.0132 | 0.6144 ± 0.0990 |
| 5 | 0.9522 ± 0.0044 | 0.0376 ± 0.0058 | 0.4904 ± 0.1106 |
| 10 | 0.9525 ± 0.0033 | 0.0429 ± 0.0128 | 0.5171 ± 0.1209 |
| 15 | 0.9496 ± 0.0033 | 0.0542 ± 0.0255 | 0.5233 ± 0.1256 |
| 20 | 0.9529 ± 0.0036 | 0.0413 ± 0.0143 | 0.5553 ± 0.0872 |
| 40 | 0.9496 ± 0.0024 | 0.0578 ± 0.0183 | 0.4935 ± 0.0961 |
| 60 | 0.9524 ± 0.0030 | 0.0421 ± 0.0076 | 0.5587 ± 0.0969 |
| 80 | 0.9507 ± 0.0031 | 0.0416 ± 0.0080 | 0.5147 ± 0.0688 |
| 100 | 0.9514 ± 0.0038 | 0.0457 ± 0.0082 | 0.4722 ± 0.1063 |

**Table 4:** Evaluation of SpliceRead across ADASYN augmentation levels for Donor. This table reports F1 Score *±* Standard Deviation (SD), Donor CMR *±* Standard Deviation (SD), and Donor NCMR *±* Standard Deviation (SD) across folds for models trained with increasing levels of synthetic non-canonical data.

| Augmentation Level | F1 Score | Donor CMR | Donor NCMR |
| --- | --- | --- | --- |
| 0 | 0.9503 ± 0.0012 | 0.0415 ± 0.0110 | 0.5034 ± 0.0415 |
| 5 | 0.9522 ± 0.0044 | 0.0335 ± 0.0060 | 0.4378 ± 0.0412 |
| 10 | 0.9525 ± 0.0033 | 0.0296 ± 0.0065 | 0.4518 ± 0.0315 |
| 15 | 0.9496 ± 0.0033 | 0.0365 ± 0.0095 | 0.4553 ± 0.0373 |
| 20 | 0.9529 ± 0.0036 | 0.0306 ± 0.0054 | 0.4540 ± 0.0466 |
| 40 | 0.9496 ± 0.0024 | 0.0404 ± 0.0124 | 0.4549 ± 0.0198 |
| 60 | 0.9524 ± 0.0030 | 0.0362 ± 0.0156 | 0.4522 ± 0.0488 |
| 80 | 0.9507 ± 0.0031 | 0.0302 ± 0.0053 | 0.4466 ± 0.0378 |
| 100 | 0.9514 ± 0.0038 | 0.0347 ± 0.0085 | 0.4651 ± 0.0903 |

### 3.2 SpliceRead Overview

To understand the impact of data augmentation, we performed five-fold cross-validation across a range of ADASYN augmentation levels and selected the best model based on its F1 score for both canonical and non-canonical splice site prediction (Tables 3 and 4). While our initial assumption was that increasing the augmentation level would lead to a gradual and consistent improvement in performance, the results instead reveal a non-linear trend. For example, in the acceptor case (Table 3), the F1 score increases from **0.9503** at 0% to **0.9529** at 20%, but then drops to **0.9496** at 40% before rising again to **0.9524** at 60%. A similar fluctuation is observed for donor predictions (Table 4), where the F1 score decreases from **0.9529** at 20% to **0.9496** at 40% and then recovers to **0.9524** at 60%.

This instability demonstrates that augmentation does not yield a consistent improvement, and that excessive synthetic sampling may introduce noisy or redundant examples that temporarily reduce model performance. Therefore, rather than assuming that “more data always helps,” we focused on augmentation levels below 20%, where the gains were both consistent and computationally efficient. Detailed analysis from 0 to 100% augmentation is provided in Supplementary material(Figure S3).To validate this decision, we computed analogous performance metrics using Borderline-SMOTE. Trends for ADASYN are shown across varying augmentation levels (Figure 2), while the corresponding Borderline-SMOTE results are provided in Supplementary Figure S4.

**Fig. 2:**
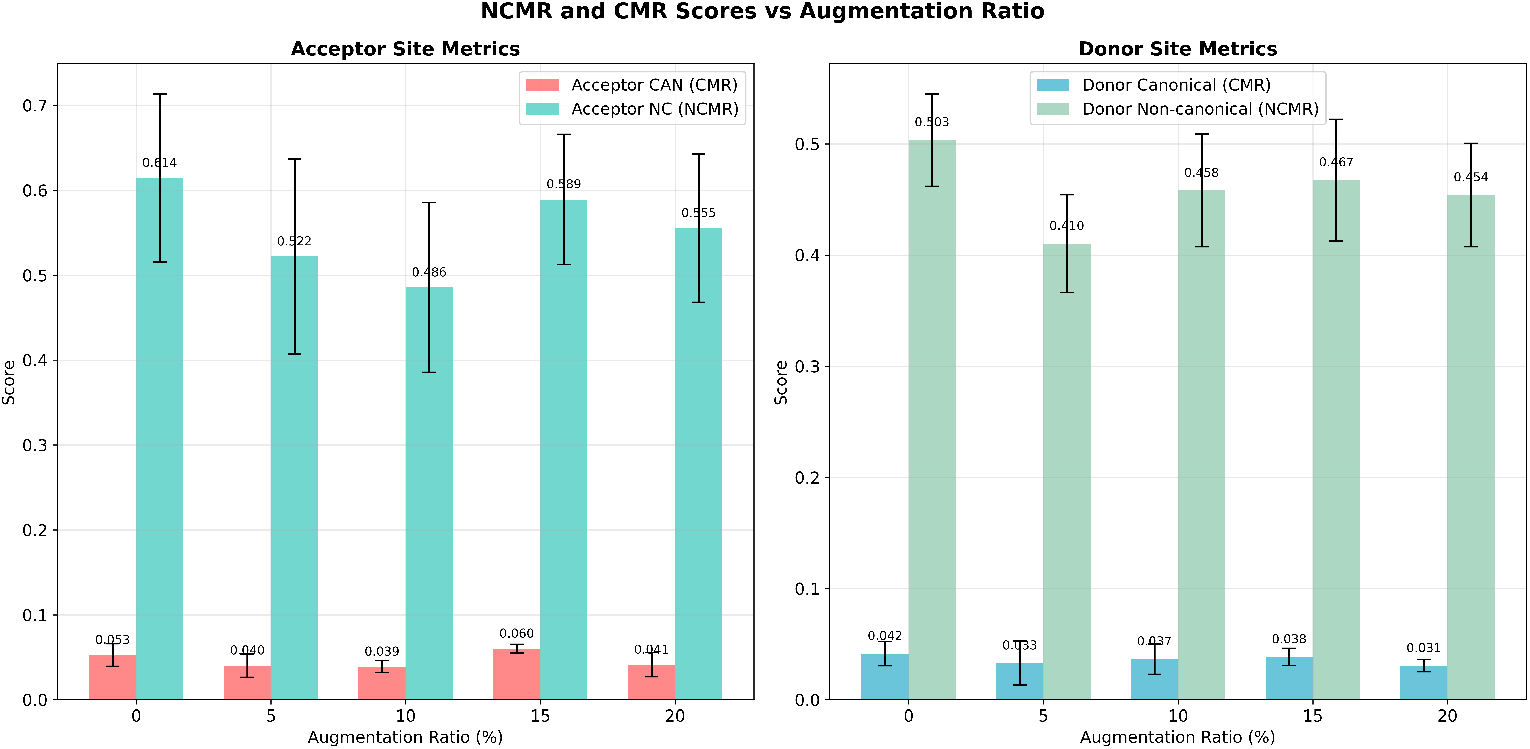
Effect of ADASYN augmentation (0–20%) on canonical (CMR) and non-canonical (NCMR) error rates for acceptor and donor splice sites

Following a detailed examination of CMR and NCMR trends (Figures 2 and Supplementary Figure S4) and guided by Occam’s Razor principle, which favors simpler solutions when performance is comparable, we selected the model trained with 5% augmented data for final benchmarking and validation, also considering that this augmentation had the lowest NCMR scores, mostly for both acceptor and donor datasets. This choice reflects the insight that minimal, targeted augmentation yielding strong predictive accuracy is preferable over a more complex model trained with excessive synthetic data. Incorporating 5% synthetic sequences improved class balance and reduced bias toward majority classes, enabling better generalization. Notably, augmentation improved F1 Score, CMR, and NCMR for both acceptor(Tables 3 and donor sites 4), demonstrating enhanced sensitivity to non-canonical splice sites, a central objective of this study.

Incorporating synthetic sequences at 5% augmentation level into the training set helped balance class distributions, enabling SpliceRead to generalize more effectively and reducing its bias toward the majority classes. Furthermore, augmentation particularly benefited F1 Score, CMR, and NCMR score for both acceptor and donor sequences, indicating improved sensitivity in detecting non-canonical splice sites, an essential goal of this study.

Following cross-validation, the best-performing model was evaluated on a held-out 20% test set that remained isolated during training and hyperparameter tuning. This model was trained using input sequences of 400 and 600 nucleotides, supporting robust performance across varying genomic window sizes (Table 5).

**Table 5:** F1-score comparison of SpliceRead using 400-nt and 600-nt input sequences.

| Input Length | F1-Score (Acceptor) | F1-Score (Donor) |
| --- | --- | --- |
| 400 nt | 0.9673 | 0.9646 |
| 600 nt | 0.9549 | 0.9568 |

### 3.3 Benchmarking with State-Of-the-art Splice Site Detectiosn Algorithms

To contextualize SpliceRead’s performance, we benchmarked it against three state-of-the-art (SOTA) splice site prediction models—SpliceFinder [1], SPLAM [4], and CNNSplice [3]. All benchmarked models were trained and evaluated on the same dataset; however, SpliceRead additionally incorporated synthetic sequences during training, as data imbalance handling is a default feature of our algorithm. Additional details are provided in Supplementary Table 2. Table 6 presents the benchmarking results across four standard classification metrics: accuracy, precision, recall, and F1-score.

**Table 6:** Benchmarking performance of SpliceRead against state-of-the-art models: SpliceFinder, SPLAM, and CNNSplice.

| Model | Accuracy | Precision | Recall | F1-Score |
| --- | --- | --- | --- | --- |
| **SpliceRead** | **0.9598-0.9614** | **0.9601-0.9614** | **0.9598-0.9616** | **0.9598-0.9614** |
| SpliceFinder | 0.9547 | 0.9547 | 0.9544 | 0.9547 |
| SPLAM | 0.9504 | 0.9504 | 0.9504 | 0.9504 |
| CNNSplice | 0.9570 | 0.9574 | 0.9570 | 0.9570 |

Although *SpliceAI* [9] represents a foundational deep learning approach for splice site prediction, we did not include it in the benchmarking experiments because the original implementation does not provide publicly available training code or configuration for retraining on custom datasets. Consequently, evaluating SpliceAI using its pre-trained model would introduce dataset and training-regime mismatches, leading to an unfair and potentially biased comparison with models retrained under identical conditions.

SpliceRead achieves the highest scores across all four metrics—accuracy, precision, recall, and F1-score, indicating superior classification performance and robustness. The model’s balanced improvements suggest a reduced trade-off between false positives and false negatives, which is particularly valuable in genomics applications where even minor misclassifications can propagate significant downstream effects. These results validate the architectural choices behind SpliceRead, particularly the incorporation of residual connections and targeted data augmentation strategies. Together, they enable the model to outperform established baselines in both canonical and non-canonical splice site classification.

### 3.4 Non-Canonical Splice Site Error Analysis and Misclassification Patterns

We performed a detailed error analysis on both canonical and non-canonical splice site predictions using the CMR and NCMR metrics. As shown in Table 7, SpliceRead consistently reduced both types of errors compared to benchmark models. For models like SPLAM, SpliceFinder, and CNNSplice, we observed minor run-to-run variations in NCMR and CMR due to the inherent stochasticity of deep learning training. Nonetheless, the trends remained consistent across experiments. These results suggest that SpliceRead exhibits a reduced bias toward canonical motifs and an improved capacity to correctly identify rare, error-prone non-canonical sequences.

**Table 7:** Classwise Canonical Misclassification Rate (CMR) and Non-Canonical Misclassification Rate (NCMR) for donor and acceptor splice sites across four benchmark models.

| Model | Acceptor CMR | Acceptor NCMR | Donor CMR | Donor NCMR |
| --- | --- | --- | --- | --- |
| **SpliceRead** | **0.0240-0.0299** | **0.5517-0.6207** | **0.0200-0.0289** | **0.3333-0.3750** |
| SPLAM | 0.0371 | 0.6876 | 0.0368 | 0.4917 |
| SpliceFinder | 0.0371 | 0.6551 | 0.0246 | 0.4792 |
| CNNSplice | 0.0425 | 0.6551 | 0.0251 | 0.4791 |

From Table 7, we observe that although SPLAM and SpliceFinder achieve similarly low CMR values on canonical splice sites, SPLAM’s much higher NCMR values for acceptor donor sites, respectively, account for its lower rankings in Table 6. Because non-canonical splice sites represent the minority class, misclassifications in this category disproportionately reduce Recall and F1-Score. Consequently, even modest reductions in NCMR lead to noticeable improvements in overall performance, which explains why SpliceFinder ranks above SPLAM, and why SpliceRead, with the lowest NCMR across both site types, achieves the strongest results across all metrics.

To further account for the extreme class imbalance inherent to non-canonical splice site prediction, we additionally report class-wise, one-vs-rest AUC-ROC and AUC-PR metrics for acceptor and donor sites separately (Table 8). While ROC-AUC captures the model’s threshold-independent ranking ability, AUC-PR provides a more informative measure under severe imbalance, where random baseline performance is proportional to the prevalence of non-canonical sites. Notably, despite numerically modest absolute values, the observed AUC-PR scores substantially exceed their respective random baselines, indicating meaningful enrichment of non-canonical splice sites. Together with NCMR and CMR, these metrics provide a granular and complementary iew of model behavior across both majority and minority splice site classes.

**Table 8:**
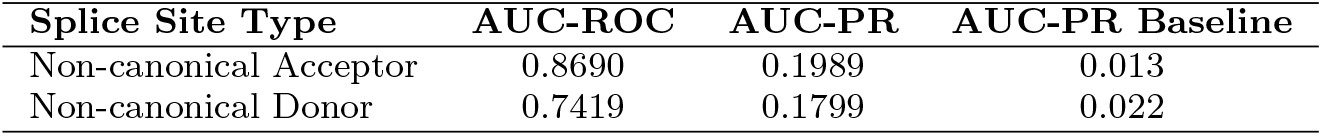
Per-class non-canonical splice site detection performance evaluated using AUC-ROC and AUC-PR. Reported AUC-PR baselines correspond to the prevalence of non-canonical sites and indicate the expected performance of a random classifier in highly imbalanced settings.

| Splice Site Type | AUC-ROC | AUC-PR | AUC-PR Baseline |
| --- | --- | --- | --- |
| Non-canonical Acceptor | 0.8690 | 0.1989 | 0.013 |
| Non-canonical Donor | 0.7419 | 0.1799 | 0.022 |

### 3.5 Cross-Species Generalization of SpliceRead versus State-of-the-art Algorithms

To evaluate the cross-species robustness of *SpliceRead*, we conducted benchmarking on two phylogenetically distinct organisms, *Danio rerio* (zebrafish) and *Drosophila melanogaster* (fruit fly), using the multi-species splice site dataset introduced in Spliceator [2]. This validation is essential to assess whether a model trained solely on species-specific sequences can generalize beyond human genomic patterns. Additional details about training these models (SPLAM, SpliceFinder, CNNSplice, and SpliceRead) are provided in Supplementary Table 2. Each model was trained on Worm data and tested on Fly and Danio data using 400-length sequences derived exclusively from the target species. No human or mixed-species data were used, ensuring performance reflects true cross-species generalization capabilities.

As shown in Figure 3, *SpliceRead* outperforms other models across all metrics and species, achieving an higher accuracy for both *Danio* and *Fly*. Its superior accuracy highlights its sensitivity in detecting splice sites across divergent genomic architectures. Notably, while CNNSplice and SPLAM also maintain competitive results. These findings validate the generalizability of SpliceRead’s architecture and underscore its potential for broader applications in comparative genomics, especially in species with limited annotated genomic data.

**Fig. 3:**
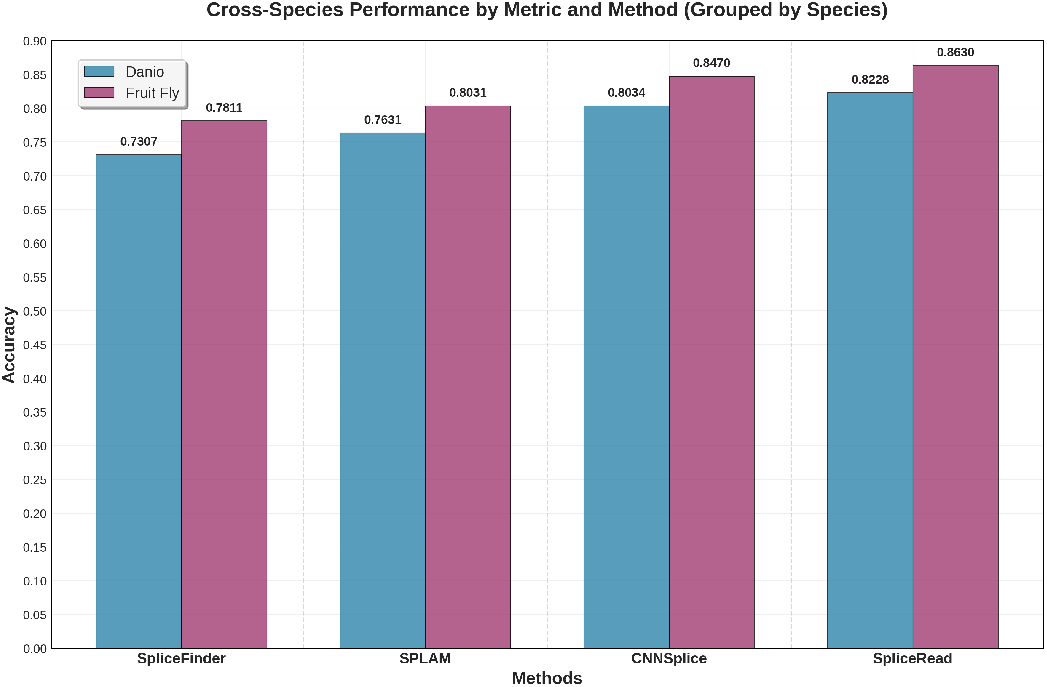
Cross-species generalization performance of SpliceRead compared to baseline models, measured by classification accuracy on two non-human species: *Danio rerio* (zebrafish) and *Drosophila melanogaster* (fruit fly).

### 3.6 SpliceRead Model Interpretability

Here, we performed two analysis to examine our model interpretability. First, we examined the sequence composition underlying splice-site regions, we first generated position weight matrix (PWM) information logos for the held-out test sets (Figure 4). Unlike attribution-based visualizations, which reflect the model’s learned decision logic, PWM logos depict the empirical nucleotide frequencies surrounding the splice junction. As expected, canonical acceptor sites exhibit a strong AG dinucleotide at positions 0/+1 together with an upstream pyrimidine-rich tract, while canonical donor sites display the conserved GT at +1/+2 followed by an extended exonic AG/G motif. In contrast, non-canonical sites show markedly weaker or more diffuse sequence enrichment, reflecting their heterogeneous biological origin.

**Fig. 4:**
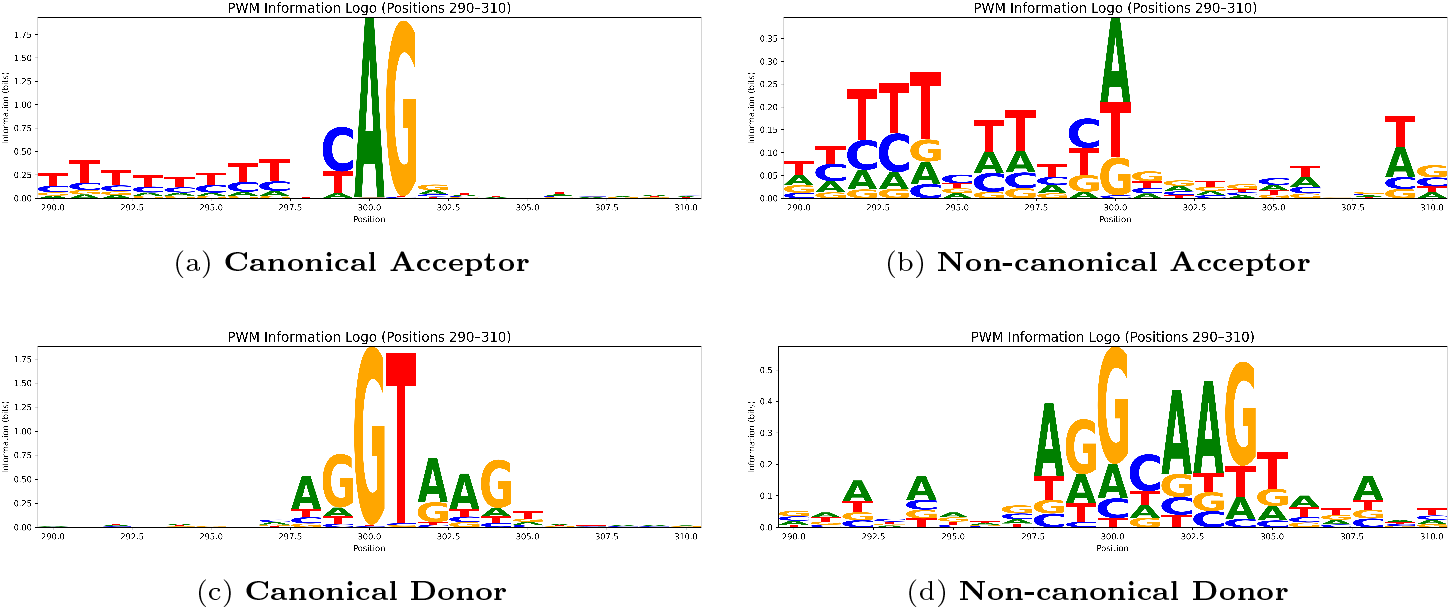
PWM information logos around the splice junction (positions 290–310). Top: acceptor sites (canonical vs. non-canonical). Bottom: donor sites (canonical vs. non-canonical). The PWM logos summarize the true nucleotide distribution in the test sequences and serve as a biological reference.

Second, to investigate which sequence features the classifier relies on for splice-site predictions, we subsequently generated position-specific signed contribution logos for the same test sets (Figure 5). We used SHAP (SHapley Additive exPlanations) [24] to compute per-base feature attributions and applied a gradient-based fallback when SHAP values were numerically negligible. For each sequence, we extracted the attribution values for the target class and aligned them with the nucleotide actually present at each position. We then averaged these signed contributions across all test samples, yielding a contribution-weighted matrix (CWM) in which positive values indicate bases that increase the model’s confidence, while negative values indicate inhibitory evidence. These aggregated attributions were visualized as signed contribution logos. When SHAP occlusion values were numerically indistinguishable from zero, we used gradient × input as an attribution fallback; this affected only the magnitude, not the location, of the dominant motifs.

**Fig. 5:**
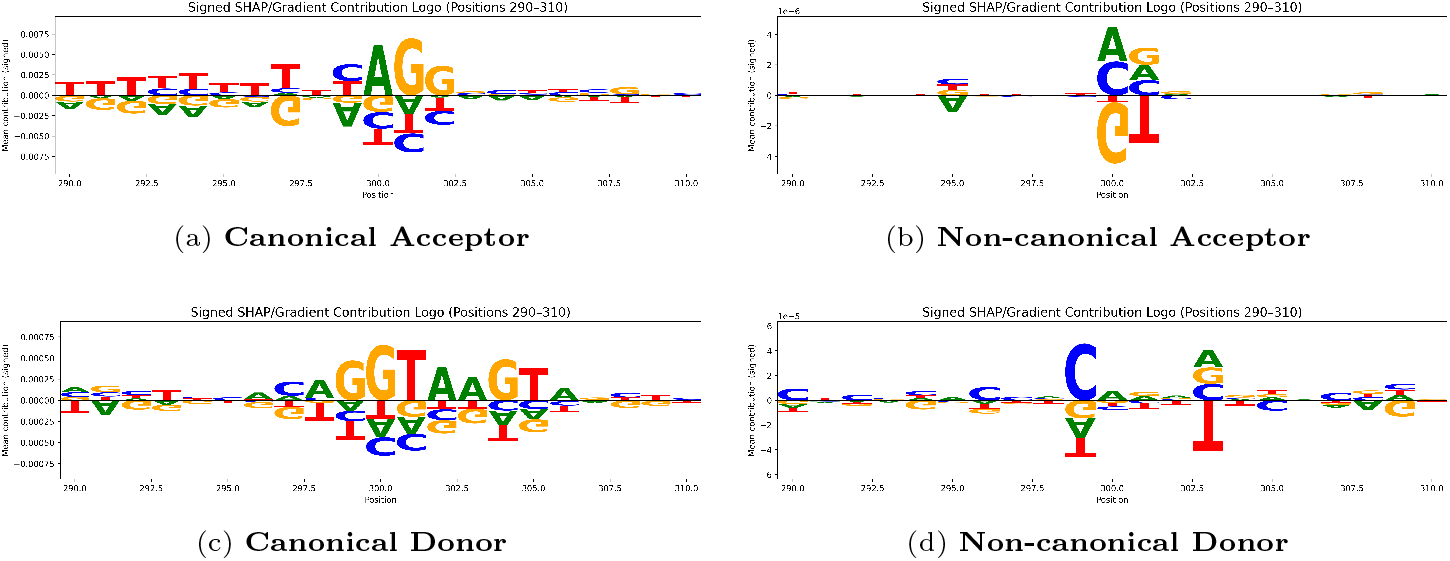
Signed contribution logos (SHAP/gradient; centered at zero) around the splice junction (positions 290–310). Top row: acceptor sites. Bottom row: donor sites. Positive letters indicate bases that increase the target logit; negative letters decrease it.

In the acceptor set, the model assigns a strong positive weight to the canonical AG dinucleotide at the splice junction and to the upstream pyrimidine tract, mirroring known splice-acceptor biology (Figure 5a). In the donor set, the model correctly recovers the canonical GT at positions +1/+2 and flanking exonic biases. The absence of spurious or off-motif signals suggests that the classifier has learned genuine splice motifs rather than dataset artifacts, providing mechanistic plausibility for its redictions (Figure 5c).

Comparing these PWMs with the signed SHAP-based attributions reveals that SpliceRead assigns high positive importance to the same canonical motifs that dominate the empirical sequence distribution. This confirms that the model amplifies genuine splice signals rather than relying on spurious features.

For non-canonical splice sites, both the PWM (Figure 4b, 4d) and attribution logos show weaker motif strength (Figure 5b, 5d), suggesting their lower predictability stems from biological sequence variability rather than a model shortcoming. Interestingly, this lack of dominant patterns in the non-canonical sites highlights the inherent volatility in these signals. However, our use of synthetic data augmentation helped the model learn more discriminative patterns even in these ambiguous regions. This suggests that the synthetic examples may have enabled the model to generalize subtle features not otherwise captured in the original imbalanced training data.

## 4 Conclusion

In this study, we introduced *SpliceRead*, a deep learning framework that enhances the prediction of canonical and non-canonical splice sites by integrating residual convolutional blocks with synthetic data augmentation. Addressing the long-standing challenge of class imbalance and sequence variability, *SpliceRead* applies ADASYN-based augmentation and systematic hyperparameter optimization to achieve robust and balanced performance. The model consistently outperformed state-of-the-art approaches across accuracy, precision, recall, and F1-score, attaining the lowest misclassification rates for both canonical and non-canonical sites. Improvements were most evident in recall, indicating stronger detection of rare non-canonical sites that often evade conventional classifiers. Moreover, *SpliceRead* maintained high cross-species ccuracy on datasets from *Danio rerio* and *Drosophila melanogaster*, demonstrating its capacity for generalization, while SHAP-based interpretability analysis verified that it captured biologically meaningful sequence motifs. Overall, *SpliceRead* provides a robust and interpretable foundation for splice site prediction and represents a significant step toward more accurate and equitable modeling of canonical and non-canonical splicing across diverse genomic systems. In the future, we plan to adapt this augmentation strategy to other state-of-the-art splice site prediction algorithms through a plug-and-play framework that enables fast integration and broad applicability. Additionally, we aim to expand our evaluation to more species and tissue types, including long-read sequencing data, to assess SpliceRead’s robustness in more biologically complex and noisy environments.

## 5 Source Code and Data Availability

The open source code of SpliceRead and a detailed Documentation is available at https://github.com/OluwadareLab/SpliceRead. The data used in this study are available at https://doi.org/10.5281/zenodo.15538290

## 7 Acknowledgment

Not Applicable.

## 8 Authors’ contributions

OO conceived the project. ST and KS designed the algorithm. ST, KS, and RM implemented the algorithm. ST, KS, and RM performed the statistical and simulation analyses. ST, KS, and OO evaluated the results and wrote the manuscripts. All authors reviewed the manuscript.

## 9 Supplementary Information

Table S1 - S2 and Figure S1 - S4

