## Supplemental Document for "SpliceRead: Improving Canonical and Non-Canonical Splice Site Prediction with Residual Blocks and Synthetic Data Augmentation"

### 1 Supplementary Materials

**Table S1:** F1-score comparison across different combinations of residual blocks, filter counts, kernel sizes (K), and pooling sizes (P). The highlighted row indicates the configuration used in SpliceRead.

| Res Blocks | Filters | K=3, P=1 | K=3, P=2 | K=5, P=1 | K=5, P=2 | K=9, P=1 / P=2 |
|---|---|---|---|---|---|---|
| 1 | 50 | 0.792 | 0.801 | 0.804 | 0.809 | 0.812 / 0.816 |
| 1 | 100 | 0.798 | 0.806 | 0.811 | 0.818 | 0.821 / 0.827 |
| 1 | 150 | 0.794 | 0.803 | 0.808 | 0.815 | 0.818 / 0.825 |
| **3** | **50** | **0.823** | **0.834** | **0.841** | **0.851** | **0.859 / 0.866** |
| 3 | 100 | 0.828 | 0.838 | 0.846 | 0.855 | 0.861 / 0.865 |
| 3 | 150 | 0.825 | 0.835 | 0.844 | 0.852 | 0.858 / 0.864 |
| 5 | 50 | 0.814 | 0.822 | 0.832 | 0.841 | 0.848 / 0.853 |
| 5 | 100 | 0.820 | 0.829 | 0.838 | 0.847 | 0.852 / 0.857 |
| 5 | 150 | 0.817 | 0.826 | 0.836 | 0.843 | 0.849 / 0.854 |

**Table S2:** Training epochs used for each model during benchmarking. SPLAM was trained for additional epochs (70 instead of 50) because its performance had not converged at 50 epochs, whereas the other models reached stable results within 50 epochs.

| Model | Epochs |
|---|---|
| SpliceRead | 50-80 |
| SpliceFinder | 50 |
| SPLAM | 65 |
| CNNSplice | 50 |

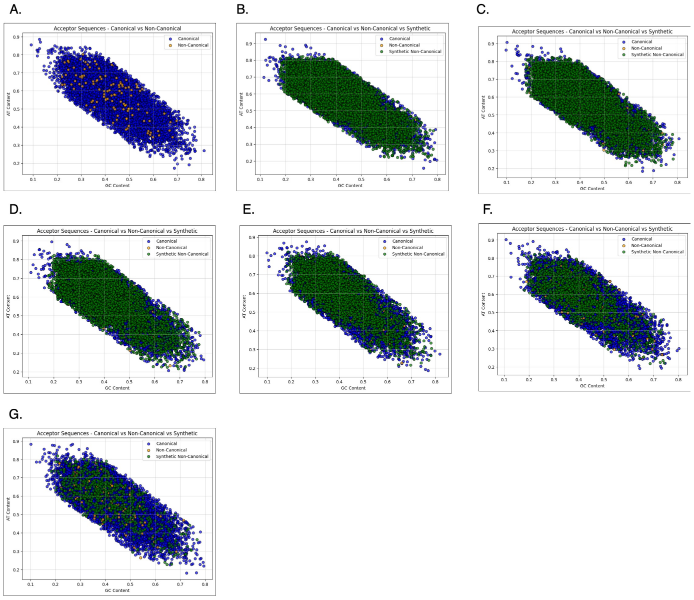


**Fig. S1:** Visualization of AT vs. GC nucleotide content in acceptor splice site sequences under progressive ADASYN-based data augmentation. Subfigure (A) shows the original distribution of canonical (blue) and non-canonical (orange) splice sites, highlighting the distinct compositional imbalance and sparsity of non-canonical sites. Subfigures (B) through (G) illustrate the introduction of synthetic non-canonical sequences (green) at decreasing augmentation levels: 100%, 80%, 60%, 40%, 20%, and 10% of the canonical class size, respectively. These synthetic samples were generated using ADASYN to address class imbalance by adaptively oversampling in regions of low density. As augmentation intensity decreases from (B) to (G), the density and coverage of synthetic non-canonical sequences decline, providing a visual representation of how augmentation levels impact feature space distribution. This figure demonstrates that synthetic sequences effectively populate underrepresented regions of the nucleotide composition space, enabling better separation and classification of non-canonical sites.

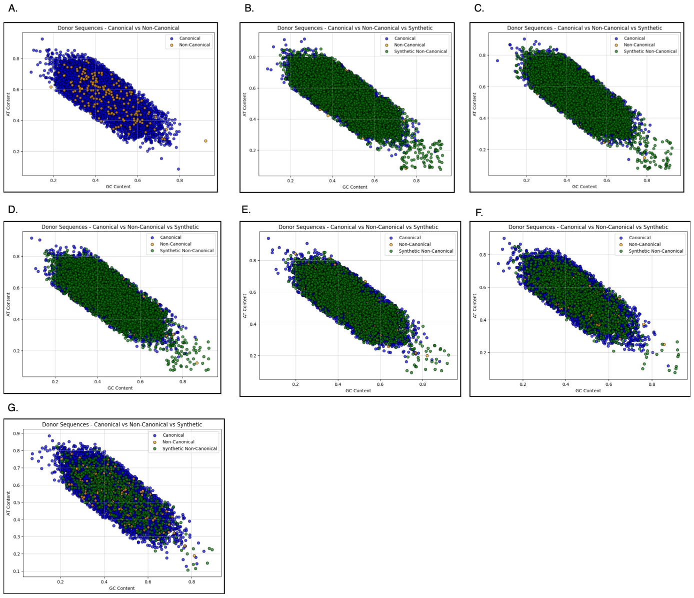


**Fig. S2:** Scatter plots illustrating AT vs. GC nucleotide content in donor splice site sequences across varying levels of ADASYN-based data augmentation. Panel (A) presents the original distribution of canonical (blue) and non-canonical (orange) donor sites, revealing a significant underrepresentation and narrower feature distribution of non-canonical sequences. Panels (B) through (G) depict the progressive incorporation of synthetic non-canonical sequences (green) generated by ADASYN at augmentation levels corresponding to 100%, 80%, 60%, 40%, 20%, and 10% of the canonical class size, respectively. Synthetic samples are generated to specifically enrich regions in the feature space where non-canonical donors are sparse, addressing class imbalance and distributional skew. As augmentation levels decrease, the spatial density of synthetic points is reduced, enabling controlled experimentation on the effect of synthetic diversity. This visualization demonstrates how augmentation reshapes the nucleotide composition landscape for donor sequences, facilitating improved model learning and generalization, particularly for rare non-canonical cases.

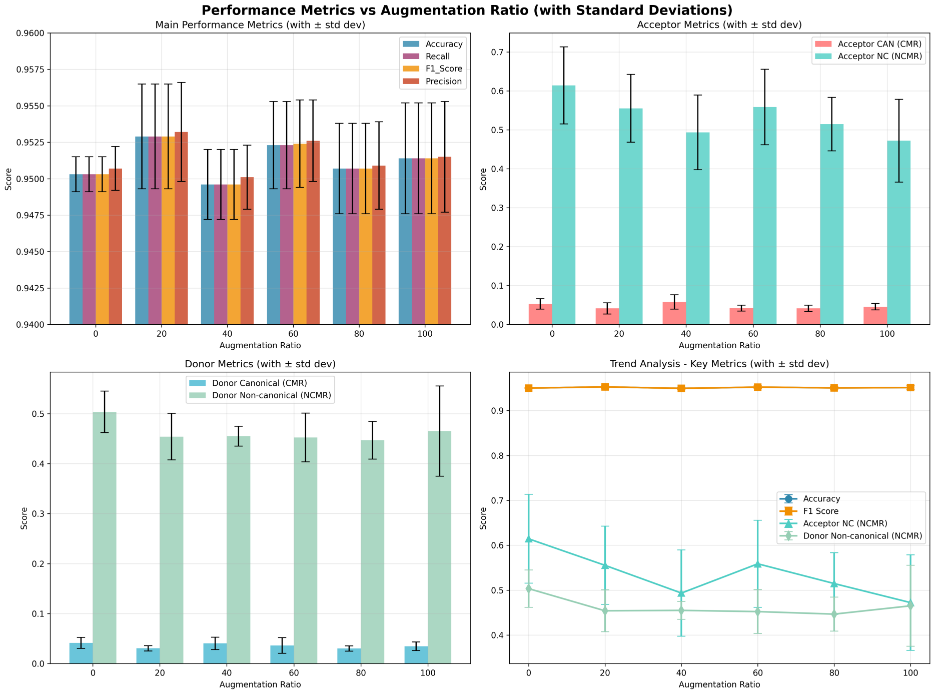


**Fig. S3: Detailed analysis of ADASYN augmentation effects across multiple evaluation metrics for SpliceRead**. Four panels summarize how performance varies as the augmentation ratio increases from 0% to 100%. (Top-left) Overall performance metrics (Accuracy, Precision, Recall, and F1-Score) remain stable across all augmentation levels, with only marginal fluctuations within the standard deviation range, indicating that augmentation does not degrade canonical classification performance. (Top-right) Acceptor-site misclassification rates show a clear reduction in NCMR between 0–60% augmentation, while CMR remains consistently low, demonstrating that synthetic oversampling primarily benefits rare, non-canonical acceptor sites. (Bottom-left) The same trend is observed for donor sites: NCMR decreases steadily up to 60%, whereas CMR stays within a narrow band below 0.05, further confirming the selective gain on minority-class predictions. (Bottom-right) The trend analysis highlights this trade-off: Accuracy and F1-Score plateau because canonical predictions dominate the dataset, while NCMR curves exhibit the most pronounced change, reinforcing that augmentation influences minority-site classification rather than global performance metrics. Collectively, these results show that ADASYN augmentation improves non-canonical site recognition without disrupting canonical accuracy, with diminishing returns beyond approximately 60% augmentation.

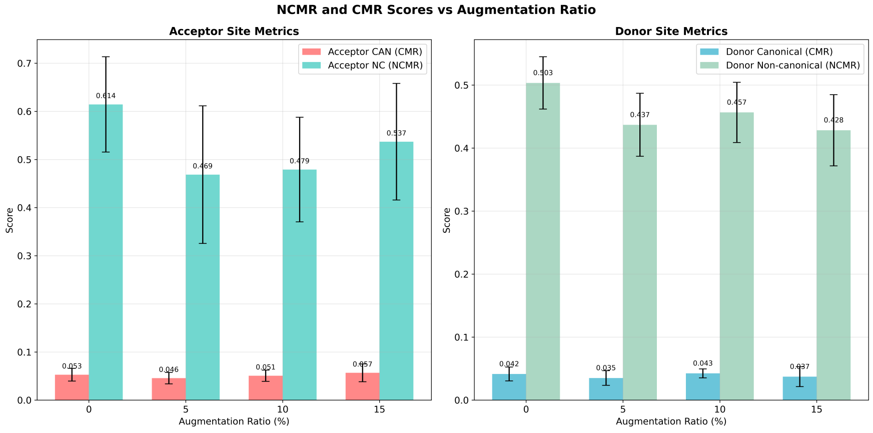


**Fig. S4: Effect of Borderline-SMOTE augmentation (0–15%) on canonical (CMR) and non-canonical (NCMR) error rates for acceptor and donor splice sites.** Mean and standard deviation are computed over five-fold cross-validation. At 5% augmentation, NCMR decreases from 0.614 to 0.469 for acceptor sites and from 0.503 to 0.437 for donor sites. In contrast, CMR changes only slightly, varying from 0.053 to 0.046 for acceptor sites and from 0.042 to 0.035 for donor sites across all augmentation levels, indicating that oversampling has a stronger effect on non-canonical splice site prediction.
